# Fitness costs of *pfhrp2* and *pfhrp3* deletions underlying diagnostic evasion in malaria parasites

**DOI:** 10.1101/2022.02.11.480127

**Authors:** Shalini Nair, Xue Li, Standwell C. Nkhoma, Tim Anderson

**Affiliations:** Disease Intervention and Prevention Program, Texas Biomedical Research Institute, San Antonio, Texas 78245; BEI Resources, ATCC, 10801 University Boulevard, Manassas, VA 20110-2209, USA

**Author notes:** **Corresponding author contact information**: Tim Anderson.

**Keywords:** Rapid Diagnostic tests (RDTs), adaptation, gene copy number, selection, *Plasmodium falciparum*

## Abstract

Rapid diagnostic tests based on detection of histidine rich proteins (HRP) are widely used for malaria diagnosis, but parasites carrying *pfhrp* deletions can evade detection and are increasing in frequency in some countries. Models aim to predict conditions under which *pfhrp2* and/or *pfhrp3* deletions will increase, but a key parameter – the fitness cost of deletions – is unknown

**Methods:** We removed *pfhrp2* and/or *pfhrp3* from a Malawian parasite clone using CRISPR/Cas9 and measured fitness costs by conducting pairwise competition experiments.

**Results:** We observed significant fitness costs of 0.087 ± 0.008 (1 s.e.) per asexual cycle for *pfhrp2* deletion and *0*.*113* ± 0.008 (1 s.e.) for the *pfhrp2*/3 double deletion, relative to the unedited progenitor parasite. The results demonstrate ∼10% reduced survival of parasites bearing deletions of these loci.

**Conclusions:** Prior modelling suggested that diagnostic selection may drive increased frequency of *pfhrp2* and *pfhrp3* deletions when fitness costs are ≤10%. Our laboratory competition experiments are consistent with costs of *pfhrp2/3* deletions lying at this critical tipping point. These results may inform future modelling efforts and help us to understand why *pfhrp2/3* deletions are increasing in some locations (Ethiopia/Eritrea) but not in others (Mekong region).

## BACKGROUND

Rapid diagnostic tests (RDTs) provide a practical way of detecting malaria (*Plasmodium falciparum*) infected individuals that can be reliably administered by public health workers with minimal training (1). These tests use a drop of blood from a finger prick that is applied to a paper strip. The sample flows laterally through a capture line of antibodies against target protein motifs, and presence off a band indicates an infection. In many endemic areas RDTs are now replacing the gold standard diagnostic approach – microscopy - which requires extensive training and validation to provide consistent results. The most widely used RDTs rely on detection of histidine rich proteins (HRP), which are among the most abundant malaria proteins and are produced across the asexual lifecycle (1). HRP-based RDTs detect products of the *pfhrp2* gene (Chromosome 8), but also show weaker detection of proteins produced by the *pfhrp3* gene (chromosome 13), which shows extensive homology (2). HRP-based RDTs show high sensitivity, because antibodies target repeat motifs in a highly expressed protein. HRP-based RDTs comprise the majority of the 345 million tests used annually, making this diagnostic tool a critical component of efforts to control and eliminate malaria from endemic areas (1, 3).

Ideally diagnostic assays should target essential genes. It is clear that *pfhrp2* and 3 genes are not essential for parasite survival, because parasites with deletions of *pfhrp2* (Dd2, D10) and *pfhrp3* (HB3) grow readily in the laboratory (4). Furthermore, parasites lacking *pfhrp2* and/or *pfhrp3* (21.6% of parasites lacking both genes) were first observed in Peru (5), and since then have been documented in several countries in South America and sub-Saharan Africa (3, 6-10). Prevalence of deletions is difficult to accurately assess, because both *pfhrp2* and *pfhrp3* are situated in highly variable telomeric regions, so PCR-based tests are prone to false negatives (2, 11). As parasites bearing *pfhrp2* and *pfhrp3* deletions evade detection by HRP-based RDTs, there are concerns that selection for diagnostic evasion could drive increases in frequencies of HRP-deleted parasites. Consistent with this, parasites bearing *pfhrp2* and *pfhrp3* deletions are now commonly found in the horn of Africa (Eritrea, Ethiopia) (3, 10) and there are strong signals of selection surrounding the HRP2 deletion, indicative of recent positive selection for this deletion (3).

Attempts to model how HRP-based diagnosis of malaria can drive increase in frequency of *pfhrp2* and *pfhrp3* deletions can be useful to (i) to understand why deletions are at high frequencies in some countries, but not others, and (ii) to prioritize countries where surveillance of deletion frequency is warranted (12, 13). Current models suggest that low prevalence and a high proportion of infected people seeking treatment are the most important drivers. However, one important parameter that is currently unknown is the fitness cost of deletions: high costs will retard rate of spread of deletions (12). Current models suggest that *pfhrp2* and *pfhrp3* deletions are unlikely to increase in frequency unless relative fitness of deletion mutants is >90% that of wildtype parasites (i.e. fitness costs < 0.1)

This project was designed to determine fitness costs of *pfhrp2* and *pfhrp3* deletions. We generated parasite clones with *pfhrp2* and *pfhrp2*/3 double deletions from a recently isolated Malawian parasite clone using CRISPR/Cas9 methods, and then measured fitness of blood stage parasites using competitive growth assays, and amplicon sequencing assays to determine the outcome of competition. Our results reveal fitness costs of 0.09 for *pfhrp2* deletions and 0.11 for the *pfhrp2*/3 double deletion, which is broadly consistent with model sensitivity analyses.

## METHODS

### Parasites

We used a parasite isolate (LA476) collected from a Malawian patient in 2008 as part of a cross-sectional study approved by the College of Medicine Research and Ethics Committee, University of Malawi. *Pfhrp2* and *pfhrp3* are located in telomeric regions of chr 8 and 13 and flanking regions show extensive sequence variation and rearrangements (2). We therefore sequenced the LA476-1 genome to ensure that guide RNAs and flanking sequences could be designed to accurately match this particular parasite isolate.

### CRISPR/Cas9 methods for generating deletions

We designed a plasmid for CRISPR/Cas9 deletion of *pfhrp2* and *pfhrp3* using CRISPR/Cas9 approaches (Fig. 1A). The guide RNAs were deigned to target positions in the LA476-1 genome (coordinates based on homology with 3D7 genome, PlasmoDB release 46) flanking *pfhrp2* and *pfhrp3*. Plasmids were constructed by Genscript, and homology arms and gRNA sequences were checked using Sanger sequencing. The homology region consists of 935-1031 bp flanking regions surrounding the region targeted for deletion. We designed sgRNA sequences (Table 1) at the 5’, 3’ or mid regions of the genes and used Cas-OFFinder software to search for potential off target sites.

**Table 1.**
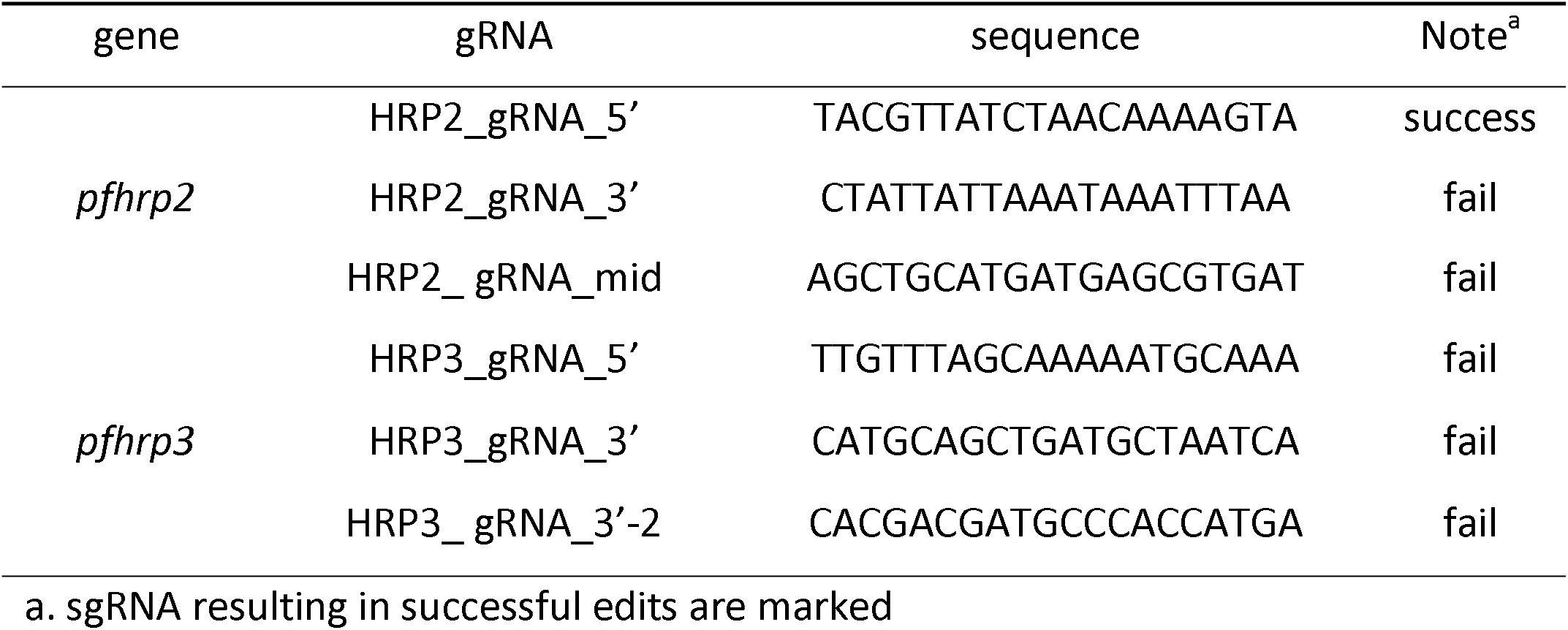
Design of sgRNA.

**Figure 1.**
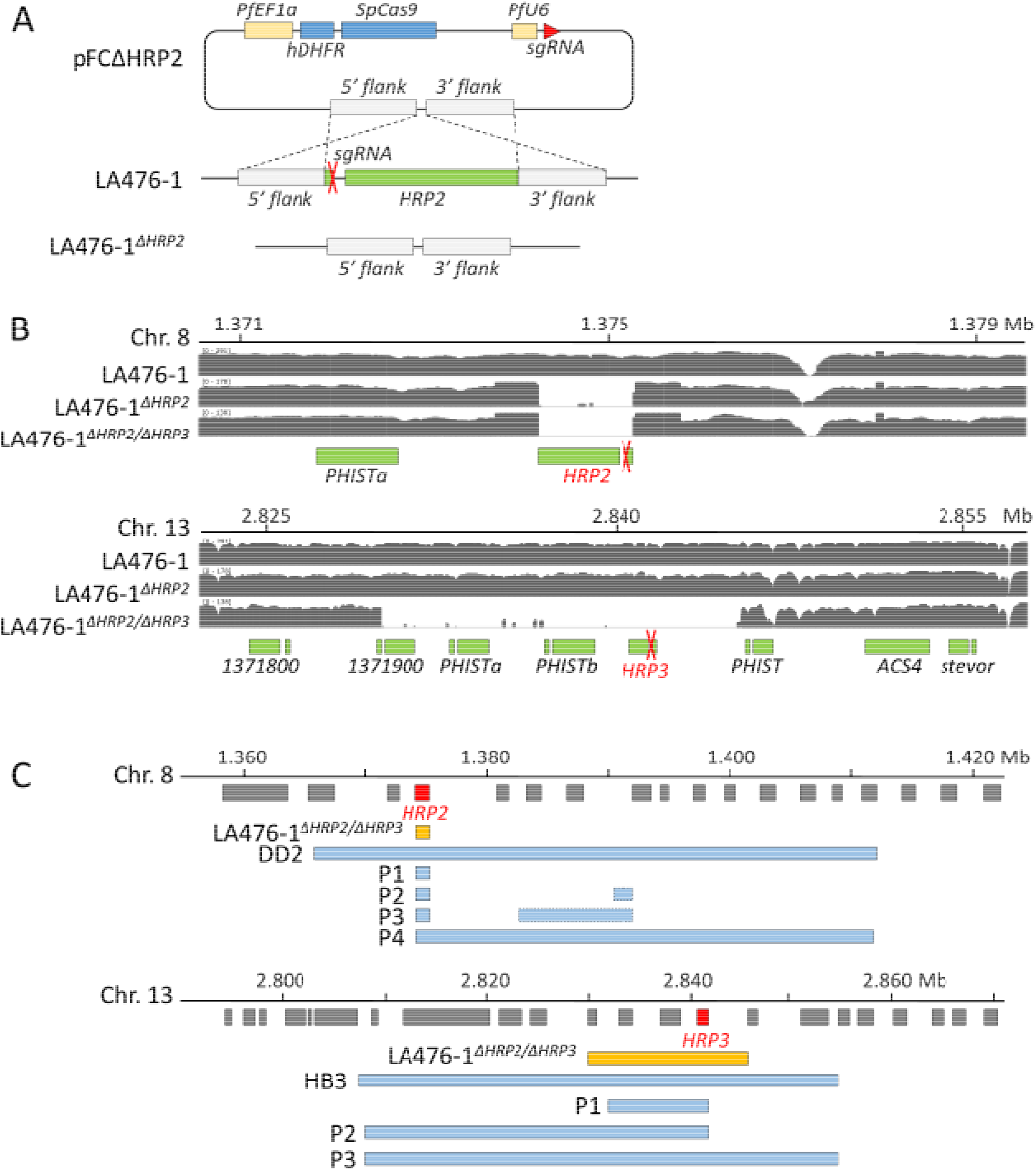
CRISPR generated *pfhrp2* and *pfhrp3* deletions. A, strategy to generate LA476-1^Δ*HRP2*^. LA476-1^Δ*HRP2/*Δ*HRP3*^ was generated in the same manner. CRISPR/Cas9 modification plasmids were generated by adding sgRNA and homology arms flanking the upstream and downstream regions of *pfhrp2* or *pfhrp3* to the pFC plasmid previously reported by Goswami et al (33). B, IGV Plots showing *pfhrp2* and *pfhrp3* deletions for parasites generated. sgRNA target loci were labeled with red crosses. C, schematic comparisons of deletions in field isolates and CRISPR generated deletions. DD2 and HB3 are lab strains that know to have HRP2 (DD2) or HRP3 (HB3) deletions. P1-4 in Chr.8 panel and P1-3 in Chr.13 panel indicate field samples from Ethiopia grouped by *pfhrp2* or *pfhrp3* deletion structural profile (3).

We transfected ring stage parasites with 100 ng plasmid DNA, and selected successful transfectants by treatment with 5 nM WR99210 (Jacobus Pharmaceuticals, Princeton, NJ) for 6 days and we recovered parasites after ∼3 weeks. To determine if recovered parasites contained the deletions expected, we conducted PCR assays using primers as showed in Table 2. We cloned parasites from successful transfection experiments. Recovered parasites were then genome sequenced (i) to define the breakpoints of deletions, (ii) to identify any off-target edits elsewhere in the genome.

**Table 2.**
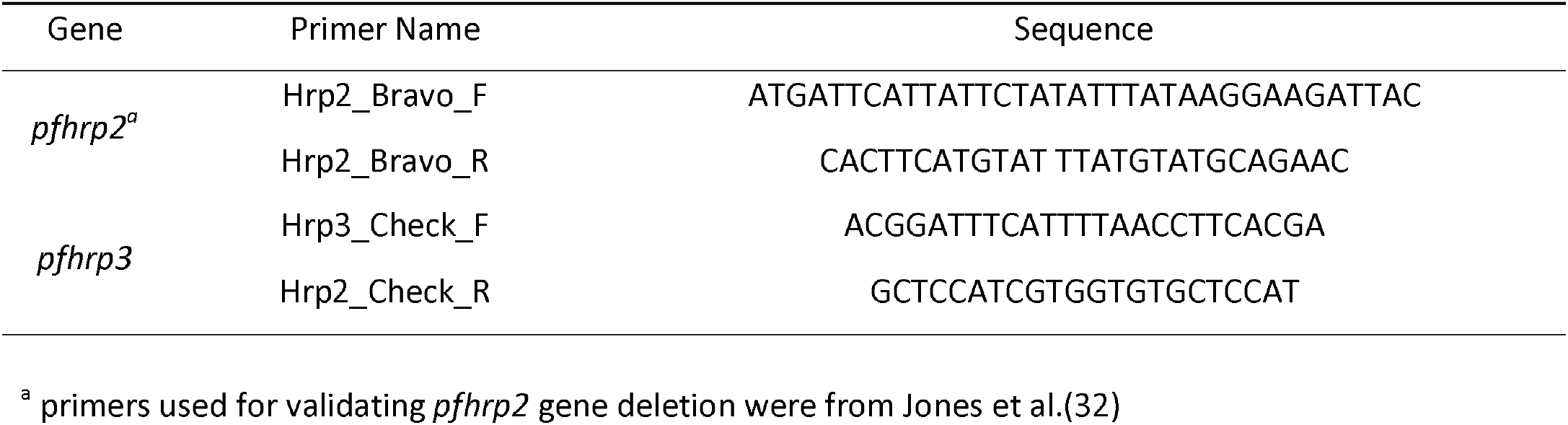
Primers used to validate *pfhrp2* or *pfhrp3* deletions.

### Measurement of fitness costs

We conducted head-to-head competition experiments to determine the impact of *pfhrp* deletions on parasite fitness in asexual culture. To do this we competed LA476 and each of our *pfhrp* deletion mutants against a common competitor (NHP4302, CRISPR edited parasite with 2 synonymous mutations at kelch13 gene locus) (14). All competition experiments were conducted with 6 replicates in 6 well plates. We synchronized parasites to 80% schizont stages by purifications through MACS purification columns (Miltenyi Biotec): after overnight growth, we plated ring stage parasites at 50% frequency (0.1% parasitemia) to initiate experiments. We maintained cultures at 0.1-4 parasitemia, and collected 80 ul aliquots every 4 days to monitor allele frequency change. We conducted these experiments using complete media consisting of RPMI 1640 supplemented with 2 mM L-glutamine, 25 mM HEPES, and 50 g/liter gentamicin, 0.1M hypoxanthine, and 0.4% AlbuMAX as a serum source. Parasites were grown at 2% hematocrit and maintained at 37°C with 5% O2, 5% CO2, and 90% N2.

We amplified a 249bp sequence region of kelch13, spanning the synonymous mutations present in NHP4302, to determine frequencies of competing parasites within mixtures. We then sequenced amplicons to high read depth on an Illumina MiSeq run to quantify allele frequencies using methodology described previously (14).

### Statistical analysis

We plotted the natural log of the parasite ratio (frequency of clone A/Frequency of clone B) against time. Outliers were removed upon Cook’s Distance (15) with a cut-off of 4/n. The slope of the best fit linear model provides the selection coefficient (*s*), a measure of relative fitness of the competing parasites. We used the R package *metaphor* (Random-Effects Model) to compare selection coefficients resulting from *pfhrp* deletions

### Data availability

Raw sequencing data have been submitted to the NABI Sequence Read Archive (SRA) (https://www.ncbi.nlm.nih.gov/sra) under project accession number PRJNA798076.

## RESULTS

### Generation of deletion mutants

We generated both the LA476 ^Δhrp2^ and LA476 ^Δhrp2/hrp3^ deletion mutants. However, we were unable to generate the LA476 ^Δhrp3^ despite 3 different attempts using different sgRNA (Table 2). The breakpoints of *pfhrp2* deletion were the same for both the LA476 ^Δhrp2^ and LA476 ^Δhrp2/hrp3^ parasites, and as expected from the CRISPR/Cas9 design. The deletion spanned 1061bp (1,374,237-1,375,298 bp) on chr 8 and contains only *pfhrp2*. The *pfhrp3* deletion was inadvertently generated using the plasmids designed to make *pfhrp2* deletions, reflecting the homology between these two genes. The *pfhrp3* deletion spans 15,566bp (2,829,897-2,845,466 bp) on chr 13 and contains three genes encoding exported proteins of unknown function - PF3D7_1371900, *PHISTa* (PF3D7_1372000) and *PHISTb* (PF3D7_1372100) in addition to *pfhrp3* (Fig 1B). For comparison, we show the gene content of naturally occurring deletions at *pfhrp2* and *pfhrp3* from Ethiopia (Fig 1C) (3). We examined genome sequences of cloned edited parasites as described in (14) for any off-target deletions or mutations relative to the original LA476 parasite. All raw data are in NCBI PRJNA798076 (https://www.ncbi.nlm.nih.gov/bioproject/PRJNA798076). We only found one SNP (Ser4026Pro) in LA476 ^Δhrp2^ that located at gene *PF3D7_1474200* (unknown function). We investigated sequences 23-bp upstream to 23-bp downstream from the position of this mutation, and compared with the guide RNA sequence. Statistical significance for the probability that these sequences are target of guide sequence were calculated using formula from Cho et al. (16). Sequences around this SNP showed no similarity to the guide sequence (*P* > 0.05), which indicates that this mutation was not caused by off-targets. Gene *PF3D7_1474200* was highly expressed in late stage gametocytes while not in other blood stage parasites (data from Malaria Cell Atlas, https://www.sanger.ac.uk/tool/mca/new/). The SNP mutation detected in this gene is unlikely to influence the quantification of parasite fitness in head-to-head experiments.

### Results of competition experiments

The common competitor outcompeted LA476, LA476 ^Δhrp2^, and LA476 ^Δhrp2/hrp3^, but the magnitude of the selection coefficient differs in each case, allowing us to determine the fitness costs of each of the deletion mutants (Table 3). We observed a fitness cost of 0.087 ± 0.008 (1 s.e.) for LA476 ^Δhrp2^ relative to LA476 (p *= 1*.*26E-06)* and *0*.*113* ± 0.008 for LA476 ^Δhrp2/hrp3^ relative to LA476 (p= *5*.*23E-08)*. The double deletion carries a significant higher fitness cost than the single deletion (p = 5.69E-04) indicating that *pfhrp3* deletion also carries a small cost (s = 0.026 ± 0.006), if we assume an additive model. We were unable to generate the LA476 ^Δhrp3^ so cannot determine whether the deletions act in an additive or epistatic manner to determine parasite fitness.

**Table 3.**
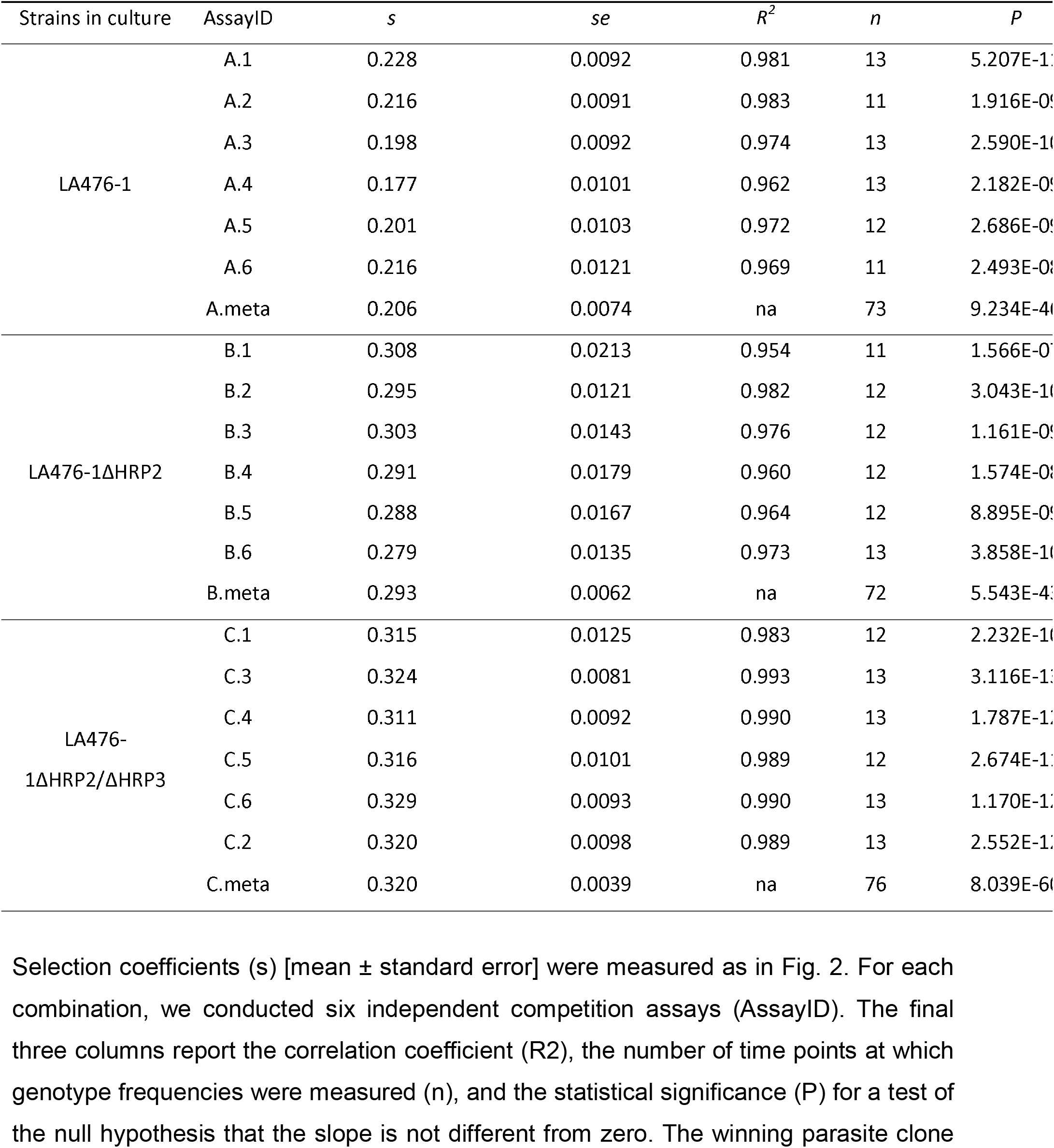
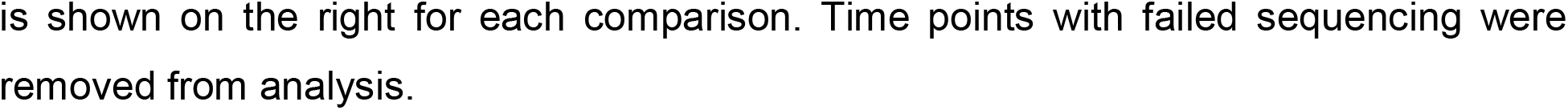
Calculation and summary for selection coefficients of all competition experiments.

## DISCUSSION

### Significant fitness costs associated with *pfhrp2* and *pfhrp3* deletions

Our results suggest a significant, fitness cost of *pfhrp2 and pfhrp3* deletions. The *pfhrp2* deletion contributes the most to the costs observed, consistent with the higher expression of this *pfhrp2 (1)*, but the *pfhrp3* gene also contributes significantly. Overall, the fitness costs are ∼0.1 – *pfhrp2/3* deleted parasites show 10% lower representation in mixtures every 48h asexual cycle. The costs are of a similar order of magnitude to those observed for many drug resistance loci in the laboratory (17).

While no other studies have directly examined fitness costs of *pfhrp2 and pfhrp3 deletion*, several other studies are informative: (i) Yang et al deleted *pfhrp2* in order to understand transcriptional consequences and function of HRP2 protein, and noted that “the growth of the parasite in the host was not affected by the deletion of *Pfhrp2* compared with the growth of the wild type”. However, these authors also determined extensive alteration in gene expression in parasites with deleted *pfhrp2*, suggesting some significant metabolic impact. We note that fitness consequences in the order of s = 0.1 may be difficult to discern without competitive growth assays. (ii) Wellems et al (18) examined the progeny from a genetic cross between HB3 (*pfhrp3* deleted) and 3D7 (no deletions), and found strong selection for progeny carrying intact *pfhrp3* in the progeny isolated, consistent with *pfhrp3* deletion, or neighboring deleted genes, carrying a fitness cost. Similarly, a cross between Dd2 (*pfhrp2* deleted) and HB3 (*pfhrp3* deleted) suggested selection against *pfhrp3*, but not against *pfhrp2* deletion (19). (iii) Zhang et al (20) conducted a piggyBAC mutagenesis study to map parasite genes that are essential and or result in reduced fitness when disrupted. Both *pfhrp2* and *pfhrp3* tolerated disruptions by multiple transposons confirming that they are non-essential (Mutagenesis Index scores (MIS) of 1 for both genes: see table S5 in (20)); furthermore, transposon disruption had little impact on competitive growth in pooled growth assays, suggesting that disabling these genes results in limited fitness costs (Mutagenesis Fitness Scores (MFS) of -1.914 (*pfhrp2*) and -1.811 (*pfhrp3*): see table S5 in (20)).

The underlying causes of slower growth in parasites bearing deletions will require improved understanding of the functions of HRP2 and HRP3 proteins. Strangely, while these genes are among the most highly expressed in the blood stage malaria parasites, we currently know rather little about their function (1). Some results suggest that HRP2 is involved in heme metabolism (21, 22): the HRP2 protein has a heme binding site (23), and deletions of *pfhrp2* results in significant under or over expression of several genes involved in hemozoin conversion (24). This feature of *pfhrp2* biology seems most likely to impact fitness in *pfhrp2* deleted parasites. Other data suggest that HRP2 is involved in capillary sequestration and cerebral malaria as *pfhrp2* alters the binding properties of erythrocytes (1, 25). Finally, *pfhrp2* may play a role in immune evasion, though suppression of B and T-lymphocyte proliferation.

### Why are deletions spreading in some locations but not others?

The balance of fitness costs, which tend to retard spread, and selection for diagnostic evasion are expected to determine spread of pfhrp2/3 deletions. Models suggest that low prevalence of malaria and high proportion of infected people seeking treatment are risk factors for spread of *pfhrp2* deletions (12). These models were missing estimates of fitness costs resulting from *pfhrp2* and *pfhrp3* deletions, but determined that deletions were unlikely to spread if fitness costs were >10%. Our estimates (9% for pfhrp2 alone, and 11% for the double deletion) fall right on this threshold. These results are consistent with the idea that deletions will start spreading only when selection for diagnostic evasion outweighs the metabolic costs resulting from deletion, and may help explain the patchy distributions of *pfhrp2* deletions. As transmission is reduced, resulting in higher numbers of symptomatic patients seeking treatment, we expect progressively more locations to reach the tipping point where benefits of *pfhrp2/3* deletions outweigh the costs, leading to an increase deletion frequency, and reduced utility of HRP2-based RDTs. Incorporating estimates of fitness costs can aid in efforts generate models to predictive prevalence of *pfhrp2/3* deletions and prioritize regions for surveillance.

In some locations, *pfhrp2* and *pfhrp3* deletions appear to have spread without selection from widespread RDT use. For example, in Amazonia *pfhrp2* and *3* deletions were first documented in 2010, and examined samples collected 2003-8, prior to widespread deployment of HRP2-RDTs (5). We suggest two related explanations for this. First, deleterious copy number variants may be more common in parasite populations with very low effective populations size (N_e_), because selection purging such alleles is weak. Consistent with such a genetic drift-based explanation, epidemic low N_e_ S. American *P. falciparum* populations carry more and larger insertions and deletions than endemic African and Asian populations with larger N_e_ (26) Second, South American parasite populations typically show highly clonal populations structure, with strong linkage disequilibrium between genes on different chromosomes (27, 28) With such population structure, deleterious mutations may spread by hitchhiking with successful multilocus genotypes. A reduction in population size, increase in genetic drift and linkage disequilibrium in populations targeted by malaria control efforts will therefore also lower the barriers against spread of mildly deleterious *pfhrp2/3* deletions.

### Limitations of study

We determined fitness during the asexual blood stage, and cannot rule out that fitness costs may be observed elsewhere during the malaria lifecycle. However, expression of *pfhrp2* and *pfhrp3* is high only in the blood stages. Neither genes are expressed in liver stage parasites, and *pfhrp2* transcripts were not detected in any mosquito stages, while *pfhrp3* shows low expressions only in activated sporozoites (29). Therefore, *pfhrp2* and *pfhrp3* seem unlikely to impact parasite fitness outside the blood stages.

We used a chloroquine sensitive African parasite for these experiments, although in South American and African countries *pfhrp2*/3 deletions have arisen in CQ-resistant parasites populations. There is some evidence that CQ-resistance and *pfhrp2* may be functionally linked (1), because *Pfhrp2* has a heme binding domain (23) and is thought to mediate conversion of heme to hemozoin in the vacuole (24). CQ resistant parasites have modified heme metabolism pathways (30), altered food vacuole pH (31), and HRP-2 mediated hemozoin conversion is pH-dependent (21). One possibility is that *pfhrp2* mediated heme -> hemozoin conversion is not utilized in CQ-resistant parasites, and that deletion of *pfhrp2*/3 will therefore have lower cost to CQ-resistant parasites. Conducting parallel *pfhrp2* and *pfhrp3* deletion experiments in CQ-R genetic background would be of considerable interest to test this hypothesis.

While our deletion on chr 8 contained only *pfhrp2*, our deletion on chr 13 contained 3 genes in addition to *pfhrp3* deletions. We cannot determine whether the fitness costs observed result from deletions of *pfhrp*3 genes, or are due to deletion of flanking genes. The extensive variation in the telomeric regions of chr 8 and chr 13 preclude precise deletion of only *pfhrp2* or *pfhrp3* genes. It is important to note that naturally occurring *pfhrp2* and 3 deletions also contain several deleted flanking genes (Fig 1C). However, the estimated cost of *pfhrp3* deletion is quite small (0.026) (Fig. 2C) suggesting that neither pfhrp3 nor flanking genes have a large impact on parasite fitness.

**Figure 2.**
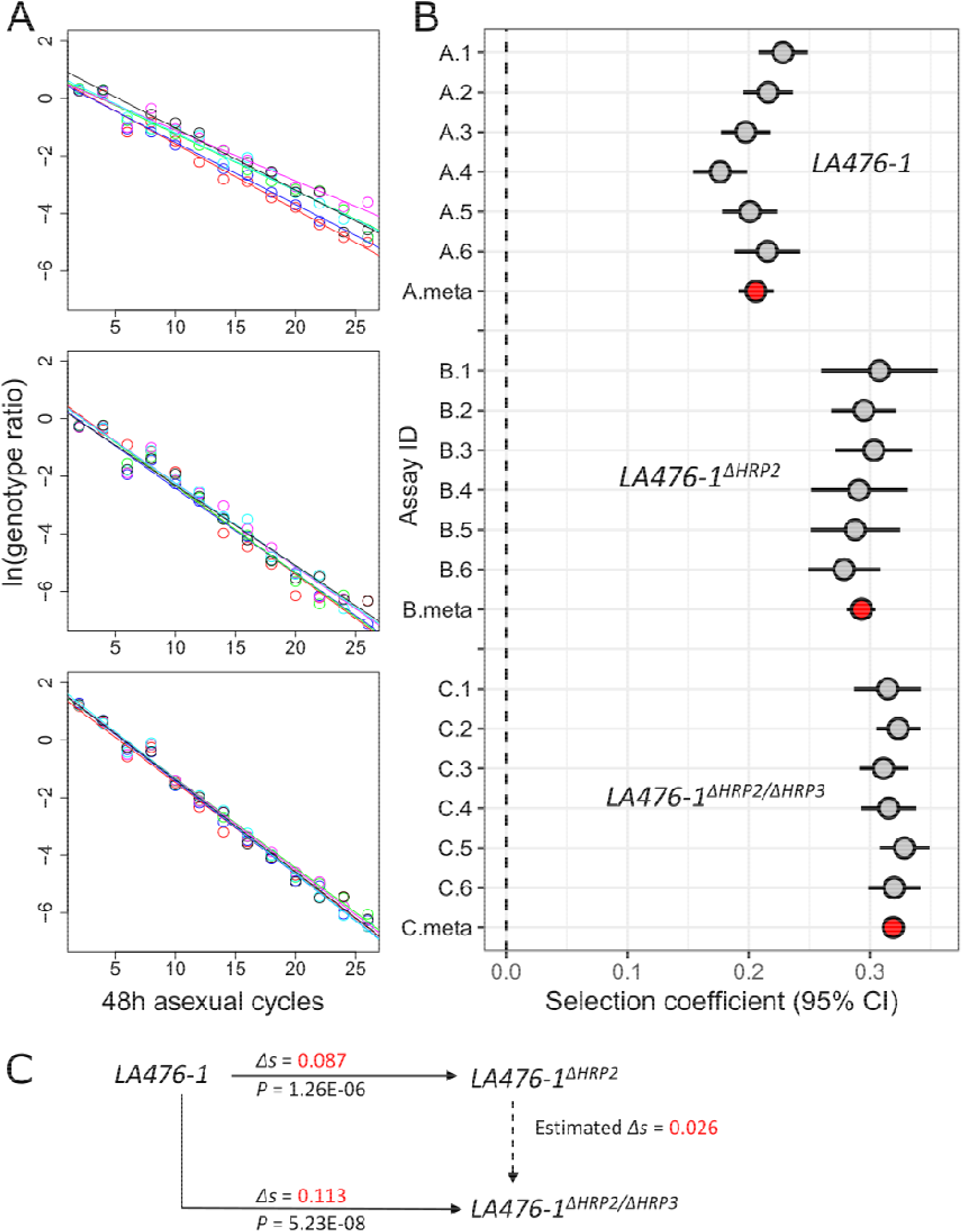
Fitness costs. A, natural log of the parasite ratio against 48-h life cycle between common competitor (NHP4026) and LA476 (top), LA476 ^Δhrp2^ (middle), and LA476 ^Δhrp2/hrp3^ (bottom). B, selection coefficients (*s*) with 95% confidence intervals from 4 sets of competition experiments. Six replicate competition experiments were conducted for each set: grey points show *s* for each replicate, while red points show meta-analysis results for each experimental comparison. C, summary of fitness differences among LA476, LA476 ^Δhrp2^, and LA476 ^Δhrp2/hrp3^.

## Acknowledgements

This work was supported by National Institutes of Health (NIH) grant R37 AI048071 (to TJCA). Work at Texas Biomedical Research Institute was conducted in facilities constructed with support from Research Facilities Improvement Program grant C06 RR013556 from the National Center for Research Resources. Collection of the parasite isolate used in this study was supported by the Gates Malaria Partnership grant to SCN. We are grateful to patients who participated in that study.

## Author contributions

S.N, X.L. and T.A. designed the experiments; S.C.N collected the parasite isolate and performed its initial characterization; S.N. conducted experiments; X.L. performed all the NGS analysis and data curation. S. N, X.L. and T.A. wrote the original manuscript.

